# drda: An R package for dose-response data analysis

**DOI:** 10.1101/2021.06.07.447323

**Authors:** Alina Malyutina, Jing Tang, Alberto Pessia

**Affiliations:** Research Program in Systems Oncology (ONCOSYS), Faculty of Medicine, University of Helsinki, Haartmaninkatu 8, 00290 Helsinki, Finland,,,, URL: helsinki.fi/en/researchgroups/network-pharmacology-for-precision-medicine/software

**Keywords:** curve fitting, dose-response, drug sensitivity, logistic function, nonlinear regression

## Abstract

Analysis of dose-response data is an important step in many scientific disciplines, including but not limited to pharmacology, toxicology, and epidemiology. The R package drda is designed to facilitate the analysis of dose-response data by implementing efficient and accurate functions with a familiar interface. With drda, it is possible to fit models by the method of least squares, perform goodness of fit tests, and conduct model selection. Compared to other similar packages, drda provides, in general, more accurate estimates in the least-squares sense. This result is achieved by a smart choice of the starting point in the optimization algorithm and by implementing the Newton method with a trust region with analytical gradients and Hessian matrices. In this article, drda is presented through the description of its methodological components and examples of its user-friendly functions. Performance is finally evaluated using a real, large-scale drug sensitivity screening dataset.

## 1. Introduction

Inferring dose-response relationships is indispensable in many scientific disciplines. In cancer research, for example, estimating the magnitude of a chemical compound effect on cancer cells holds substantial promise for clinical applications. The dose-response relationship is often modeled via a nonlinear parametric function expressed as a dose-response curve. The fitting of a curve to dose-response measurements is often achieved by choosing the parameter values that minimize the difference between the curve and the observations. Since conclusions about efficacy are based on the estimated dose-response curve, it is therefore of great importance to determine the curve parameters as accurately as possible.

Currently, there are multiple `R` packages that provide tools for the dose-response fitting, such as **drc** (Ritz, Baty, and Gerhard 2015), **nplr** (Commo and Bot 2016), and **DoseFinding** (Bornkamp, Pinheiro, and Bretz 2019). The **drc** package contains various functions for nonlinear regression analysis of biological assays. It allows the user to choose a nonlinear model for the dose-response curve fitting from a wide spectrum of sigmoid functions, which are normally used to capture the dose-response relationship as their S-shape is in line with empirical observations from experiments. The most common model is the 4-parameter generalized logistic function.

In the **drc** package, a user can specify initial model parameters to facilitate the optimization process or rely on the default starter functions. The package also enables a user to set the weights for the observations to adjust the possible variance heterogeneity in the response values. The parameter estimation procedure is achieved by the least squares method, using a maximum likelihood approach with the default assumption of normality for inferential purposes.

In contrast to **drc**, the **nplr** package focuses only on generalized logistic models and does not allow to select the data distribution. As a new feature, the package facilitates the choice of observation weights via implementing three options: residual-based, standard (or within-replicate variance-based), and general, which utilizes the fitted response values. Additionally, the package provides confidence intervals on the predicted doses and the trapezoid and the Simpson’s rule (Abramowitz and Stegun 1965, Chapter 25) to evaluate the area under the curve.

The **DoseFinding** package provides more flexibility than **drc** and **nplr**. It allows for the fitting of multiple linear and nonlinear dose-response models and to design dose-finding experiments. Similarly to **drc**, it provides several options for the data distribution, but as default it uses assumption of normality with equal variance. Compared to **drc** and **nplr**, the **DoseFinding** package utilizes a grid search as a starting point selection method in case the user did not specify its own. It also applies boundaries to parameters of a nonlinear model either specified by a user or through internal default settings.

To find the optimal parameter in a high-dimensional space, all packages apply iterative Newton methods, which are widely used numerical procedures for finding the minimum of a differentiable function (Nocedal and Wright 2006). The **drc** package directly calls the `R optim()` function that implements the Broyden-Fletcher-Goldfarb-Shanno (BFGS) method (Fletcher 2000) for unconstrained optimization, or limited-memory BFGS (L-BFGS-B), which handles simple box constraints for constrained optimization (Liu and Nocedal 1989). These two methods represent quasi-Newton methods, which are frequently used in cases when the function derivatives are not feasible or too complicated to obtain, as they utilize numerical approximations of the function’s Hessian matrix. In contrast, the **nplr** package relies on the `nlm()` function, which uses the classic Newton approach. By default, both the gradient and Hessian are approximated numerically, however the user can provide themselves the first and second analytical derivatives. The **DoseFinding** package applies different optimization routines depending on the models of choice. For sigmoid and logistic models, which have two linear and two nonlinear function parameters, the package performs numerical optimization just for nonlinear ones, while optimizing the linear parameters in every iteration of the algorithm. At its core, **DoseFinding** applies the `R nlminb()` function, using a quasi-Newton algorithm similar to the BFGS method utilized by **drc**.

While all packages have been extremely helpful with a wide range of real applications, we found that they often present inconsistent results when applied to the same data with the same logistic model. We introduce here the R package **drda**, which provides a novel and more accurate dose-response data analysis using logistic curves via: (i) applying a more advanced Newton method with a trust region; (ii) relying on analytical gradient and Hessian formulas instead of numerical approximations; (iii) establishing a smart initialization procedure to increase the chances of converging to the global solution; (iv) providing tools to compare the fitted curve against a linear model or other logistic models; (v) computing confidence intervals for the estimated parameters and for the whole dose-response curve; (vi) implementing plot functionality to compare multiple models in a user-friendly way.

The most important feature of any optimization routine remains the closeness of its solution to the true least square estimates. In case of biological assays, it depends on the ability of the fitted curve to describe the dose-response data correctly. One of the main disadvantages when it comes to numerical optimization is the possibility of converging to a local optimum instead of the correct answer we seek. This situation can easily happen when either the function is not well approximated by a quadratic shape in a neighborhood of the current candidate solution, or when the starting point is far from the global optimum (either the algorithm is not able to converge in a reasonable number of steps or it simply converges to a wrong solution). To cope with such scenarios, we implement here the Newton method with a trust region (Steihaug 1983), which has been shown to be a robust optimization technique for mitigating issues usually encountered in unconstrained optimization problems. The method is more stable than other Newton-based methods, especially for cases when it is problematic to approximate a function with a quadratic curve (Sorensen 1982). Additionally, **drda** uses a two-step initialization algorithm in order to ensure the right direction in the optimization routine. With our strategy, **drda** is able to find the true least squares estimate in problematic cases where the **drc**, **nplr**, and **DoseFinding** packages instead fail.

Once the least squares estimate is found, **drda** provides the user with routines for assessing goodness of fit and reliability of the estimates. Assuming a Gaussian distribution with equal variance for the observed data, it is possible to compare the fitted model against, for example, a flat horizontal line or a logistic model with a different number of parameters. The **drda** package provides the likelihood ratio test (LRT), the Akaike information criterion (AIC) (Akaike 1974), and the Bayesian information criterion (BIC) (Schwarz 1978) as a way to compare the goodness of fit of competing models.

The paper is organized as follows: We first describe the methodological components of **drda** in Section 2; show how the package is implemented in Section 3; include practical examples in Section 3.2; and provide a comparison of **drda** against packages **DoseFinding**, **drc**, and **nplr** using a high-throughput dose-response dataset in Section 4. We conclude the article with a summary and discussion in Section 5.

## 2. Methodological framework

### 2.1. Generalized logistic function

Package **drda** implements the generalized logistic function as the core model for fitting doseresponse data. The generalized logistic function, also known as Richards’ curve (Richards 1959), is the 5-parameter function

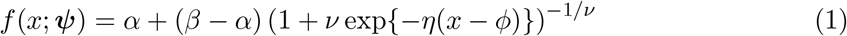

solution to the differential equation

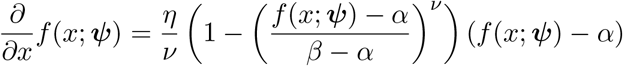

where ***ψ*** = (*α, β, η, ϕ, ν*)^*T*^.

Throughout this article, and in our package, we will use the convention *β* > *α* to avoid identifiability issues. For example, when *β* < *α*, it is always possible to modify the remaining three parameters to obtain an equivalent function. To have a sigmoidal curve, a common requirement in dose-response data analysis, we will also assume that *ν* ≥ 0. When *ν* < 0, in fact, the curve is unbounded or even complex.

Our constraints have the benefit of giving the five parameters a clear and easy interpretation: *α* is the lower horizontal asymptote of the curve, *β* is the upper horizontal asymptote of the curve, *η* is the steepness of the curve where a positive (negative) value corresponds to a monotonically increasing (decreasing) function, *ϕ* is related to the value of the function at *x* = 0, and *ν* regulates at which asymptote is the curve maximum growth. Refer to Figure 1 for a visual explanation of the five parameters.

**Figure 1:**
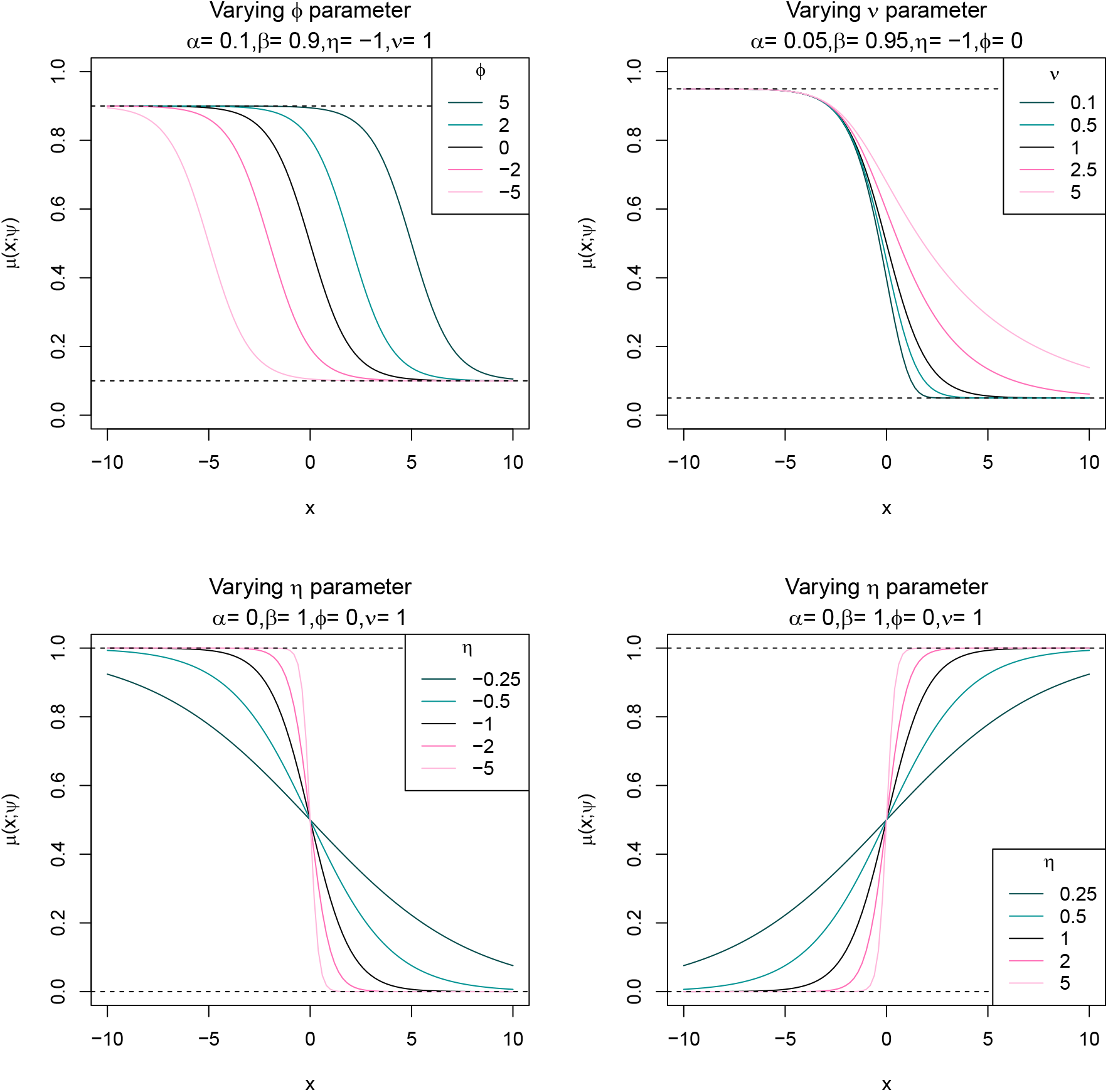
Generalized (5-parameter) logistic function with various choices of parameters.

When *ν* = 1 we obtain the *4-parameter logistic function*. If we also set *α* = 0 and *β* = 1 we obtain the *2-parameter logistic function*. When *ν* = 1 the parameter *ϕ* represents the value at which the function is equal to its midpoint, that is (*α* + *β*)/2. In such a case, as a measure of drug potency, *ϕ* is also known as the *half maximal effective log-concentration* or log-EC_50_. As a a measure of antagonist drug potency, *ϕ* is also known as the *half maximal inhibitory log-concentration* (log-IC_50_). When *ν* → 0 we obtain the *Gompertz function*, i.e.

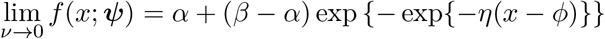

The *E_max_* model (Macdougall 2006), often found in dose-response studies, is formally equivalent to the 4-parameter logistic function. The difference between the two models is simply the parametrization of the scale used for the variable *x*. If the *E_max_* model is defined as

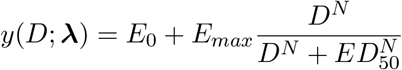

then the equivalent 4-parameter logistic function (*ν* = 1) is obtained by the transformations *D* = *e^x^*, *E*_0_ = *α*, *E_max_* = *β* – *α, N* = *η*, *ED*_50_ = *e^ϕ^*.

### 2.2. Normal nonlinear regression

For a particular dose *d_k_*(*k* = 1,…, *m*) let (*y_ki_, w_ki_*)^*T*^ represent respectively the *i*-th observed outcome and its associated positive weight. If observations have all the same importance, we simply set *w_ki_* = 1 for all *k* and *i*. We assume that each unit has expected value and variance

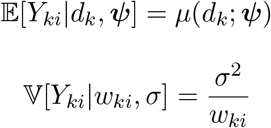

where *μ*(*d_k_; **ψ***) is a nonlinear function of the dose *d_k_* and a vector of unknown parameters ***ψ***. Parameter *σ* > 0 is instead the standard deviation common to all observations. In our package, *μ*(*d_k_*; ***ψ***) is simply the generalized logistic function (1) with the transformation *x* = log(*d_k_*).

By assuming the observations to be stochastically independent and Normally distributed, the joint log-likelihood function is

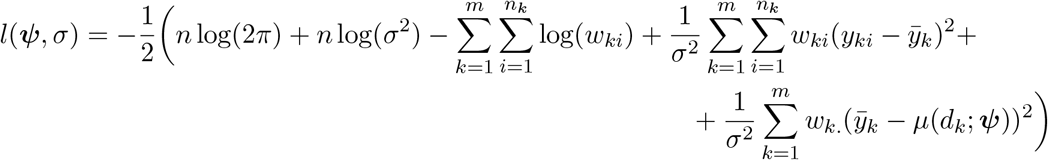

where *n_k_* is the sample size at dose *k, n* = Σ_*k*_ *n_k_* is the total sample size, 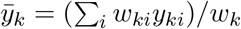 is the weighted average corresponding to dose *d_k_* and *w_k._* = Σ_*i*_ *w_ki_*. Maximum likelihood estimate 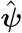 is obtained by minimizing the residual sum of squares from the means, i.e.

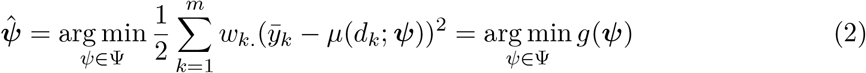

Maximum likelihood estimate of the variance is

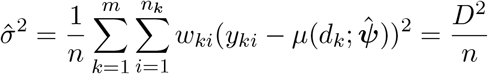

while its unbiased estimate is

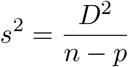

where *p* is the total number of parameters estimated from the data.

For convenience from now on we will use the simplified notation *μ_k_* to denote the function *μ*(*d_k_; **ψ***). It is important to remember that *μ_k_* will always be a function of a dose *d_k_* and a particular parameter value ***ψ***. We will also use the notation *g*^(*s*)^ and *g*^(*st*)^ to denote respectively the first- and second-order partial derivatives of function *g*(***ψ***), with respect first to *ψ_s_* and then *ψ_t_*.

Partial derivatives of the sum of squares *g*(***ψ***) are

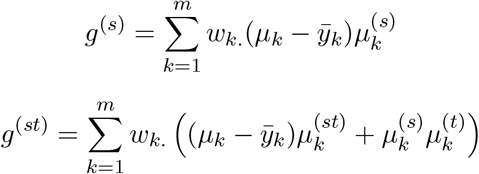

The gradient and Hessian of *g*(***ψ***) are therefore

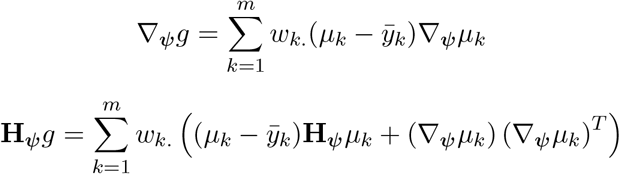

From the previous expressions we can easily retrieve the observed Fisher information matrix, which is the negative Hessian matrix of the log-likelihood evaluated at the maximum likelihood estimate, as

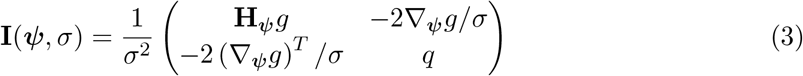

where

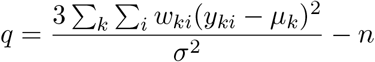

It is also worth noting that the (expected) Fisher information matrix is

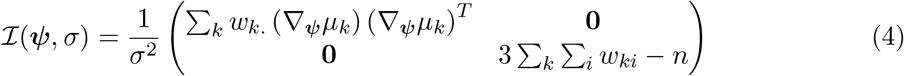

### 2.3. Optimization by Newton method with a trust region

Closed-form formula of the maximum likelihood estimate 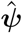, that is the solution of equation (2), is in general not available for nonlinear regression models. We can, however, try to minimize numerically the sum of squares *g*(***ψ***).

Suppose that our algorithm is at iteration *t* with current solution ***ψ***_*t*_. We want to find a new step *u* such that *g*(***ψ***_*t*_ + *u*) < *g*(***ψ***_*t*_). We start by illustrating the standard Newton method. We approximate our function by a second-order Taylor expansion, that is

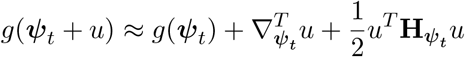

The theoretical minimum is obviously attained when the gradient with respect to *u* is zero, that is ∇*_ψ_t__* + **H***_ψ_t__ u* = 0 or 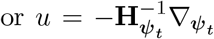. The Newton’s candidate solution for iteration *t* + 1 is often presented as

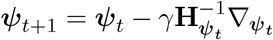

where 0 < *γ* ≤ 1 is a modifier of the step size for ensuring convergence (Armijo 1966).

When the method converges the algorithm is quadratically fast, or at least superlinear (Bonnans, Gilbert, Lemarechal, and Sagastizábal 2006): the closer *g*(***ψ***) is to a quadratic function the better its Taylor approximation, the better the algorithm convergence properties.

When the Hessian matrix is almost singular it is still possible to apply quasi-Newton methods (Luenberger and Ye 2008) to (try) avoid convergence problems. In our nonlinear regression setting, however, we might have the extra complication of an objective function far from a quadratic shape, so that the (quasi-)Newton method might fail to converge. Although this situations can be thought to be rare, they are often encountered in real applications. For example, in Figure 2 we show a problematic surface that the quasi-Newton BFGS algorithm, as implemented by the base `R` function `optim()`, is not able to properly explore.

**Figure 2:**
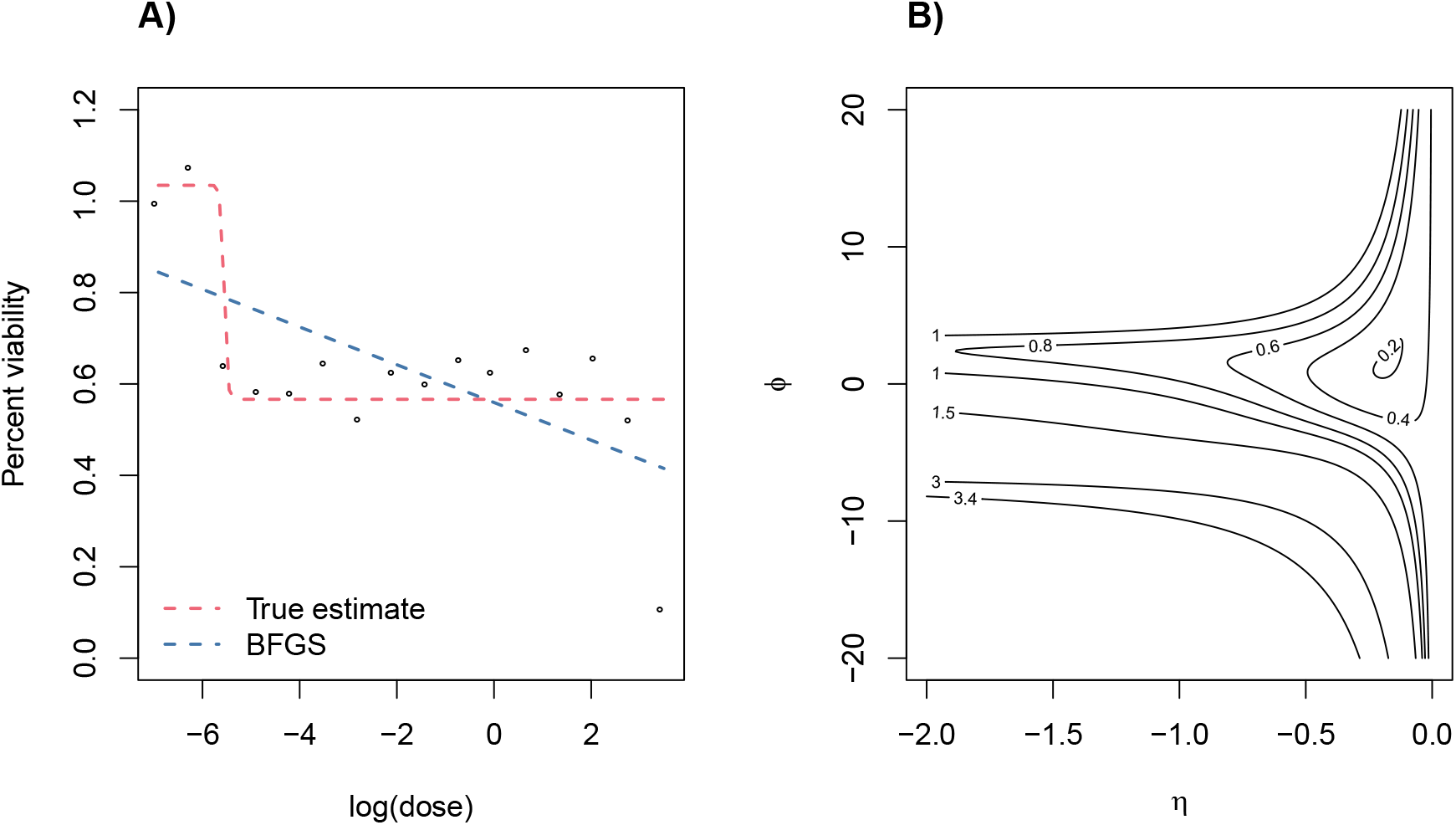
Problematic real data (cell line: BT-20, compound: BI-2536, dataset: CTRPv2) (Rees *et al*. 2016; Seashore-Ludlow *et al*. 2015; Basu *et al*. 2013)). A) 4-parameter logistic function as fitted by the BFGS algorithm. Starting point ***ψ*** = (*α, β, η, ϕ*)^*T*^ = (0, 1, −1, 0)^*T*^. B) Contour plot of the residual sum of squares *g*(***ψ***) with respect to parameters *η* and *ϕ*. Fixed parameters *α* = 0 and *β* = 1.

We will try to overcome the issues in the optimization by focusing our search only in a neighborhood of the current estimate, that is using a trust-region around the current solution ***ψ***_*t*_. The problem to solve is now

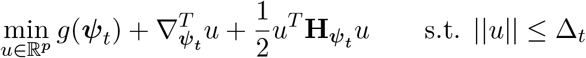

where Δ_*t*_ > 0 is the trust-region radius. Our implementation is based on the exposition of Nocedal and Wright (2006) and follows closely that of Mogensen and Riseth (2018). Briefly, at each iteration we compute the standard Newton’s step and accept the new solution if it is within the trust-region. If the Newton’s step is outside the admissible region we try an alternative step by a linear combination of the Newton’s step and the steepest descent step, with the constraint that its length is exactly equal to the radius Δ_*t*_ (dogleg method). This new alternative step is then accepted or rejected on the basis of the actual reduction in the function value. The region radius Δ_*t*+1_ for iteration *t* + 1 is adjusted according to the length and acceptance of the step just computed. For more details, we refer the reader to the extensive discussion found in Nocedal and Wright (2006).

### 2.4. Algorithm initialization

One of the major challenges in fitting nonlinear regression models is choosing a good starting point for initializing the optimization algorithm. Looking at the example in Figure 2, the choice of ***ψ***_0_ = (0, 1, −1, 0)^*T*^ made the BFGS algorithm converge to a local optimum while a global optimum might have been found if a better starting point was chosen.

First of all, we present the closed-form maximum likelihood estimates 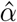 and 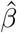 when all other parameters have been fixed. Define *h_k_* = (1 + *ν* exp(−*η*(*x_k_* – *ϕ*)))^−1/*ν*^, where *x_k_* = log(*d_k_*), and assume it to be known. Our mean function is now

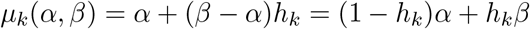

while the residual sum of squares becomes

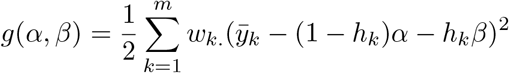

with gradient

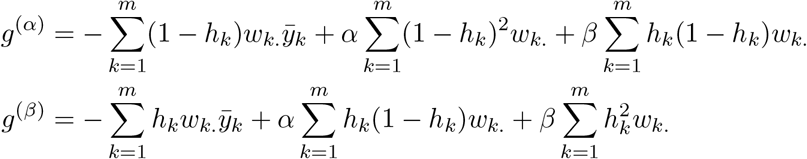

It is easy to prove that the gradient is equal to zero for

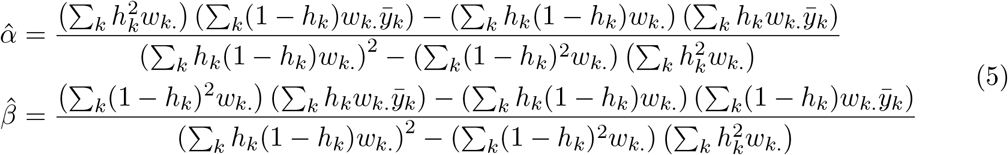

Our initialization strategy is made of two steps. The first step is done by setting *ν*_0_ = 1 and obtaining an initial guess for *η*_0_ and *ϕ*_0_, for example by choosing them at random or by evaluating the objective function on a small grid of values. We then evaluate the maximum likelihood estimates (5) and set 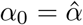 and 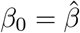. The second step is running the standard Newton method starting from ***ψ***_0_. The solution just found is then passed to our trust region implementation for further refining.

When the likelihood function is well-behaved, the standard Newton method in the second step is very fast and efficient, and most of the times will converge to the global optimum. However, when the function is problematic, we sacrifice speed for accuracy by supplying our trust region method with the local optimum found so far.

### 2.5. Statistical inference

When closed-form solutions of maximum likelihood estimates are missing, also closed-form expressions of other inferential quantities are not available. Fortunately, we can still rely on asymptotic, large sample size considerations, to obtain approximate values of quantities of interest. Obviously, the larger the sample size the better the approximation.

Using either versions (3) or (4) of the Fisher information matrix we can calculate approximate confidence intervals. In fact, we can think of the Fisher information matrix as an approximate precision matrix, so that we only have to invert the matrix and take diagonal elements as approximate variance estimates. In our package we use the observed Fisher information matrix (3) because it is shown to perform better with finite sample sizes (Efron and Hinkley 1978). As an example, an approximate confidence interval for generic parameter *ψ_j_* is

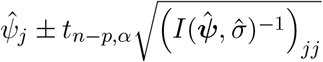

where *t_n-p,α_* is the appropriate quantile of level *α* of a Student’s *t*-distribution with *n* – *p* degrees of freedom and 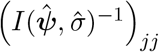 is the *j*-th element in the diagonal of the inverse observed Fisher information matrix. Using the Delta method we can compute approximate point-wise confidence intervals for the mean function

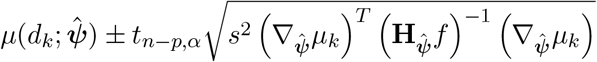

or for a new, yet to be observed, value *y*(*d*)

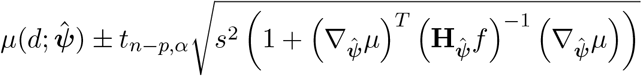

We can also construct a (conservative and approximate) confidence band over the whole mean function *μ*(·; ***ψ***) with the correction proposed by Gsteiger, Bretz, and Liu (2011)

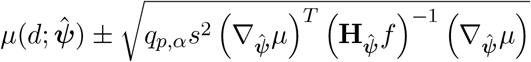

where *q_p,α_* is the appropriate quantile of level *α* of a *χ*-distribution with *p* degrees of freedom.

## 3. Using drda

### 3.1. General overview

The main function of **drda** is `drda()` with signature

~~~
drda(
  formula, data, subset, weights, na.action, mean_function = “logistic4”,
  is_log = TRUE, lower_bound = NULL, upper_bound = NULL, start = NULL,
  max_iter = 500
)
~~~

The first argument, `formula`, is a symbolic representation in the form `y ~ x` of the model to be fitted, where `y` is the vector of responses and `x` is the vector of log-doses.

`data` is an optional argument, typically a `data.frame` object, containing the variables in the model. When `data` is not specified, the variables are taken from the environment where the function is being called.

`subset` is a logical vector, or a vector of indices, specifying the portion of `data` to be used for model fitting.

`weights` is an optional argument that specifies the weights to be used for fitting. Usually weights are used in situations where observations are not equally informative, i.e. when it is known that some of the observations should have a smaller or larger impact on the fitting process. If the `weights` argument is not provided then the ordinary least squares method is applied.

`na.action` defines a function for handling `NA`s found in `data`. The default option is to use `na.omit()`, i.e. to remove all data points associated with the missing values.

`mean_function` argument specifies the model that should be estimated. In the current version of the package the argument can be any of `‘logistic5’, ‘logistic4’, ‘logistic2’`, or `‘gompertz’`. Each model is explained in detail in Section 2.1. By default, the 4-parameter logistic function is chosen.

`is_log` is a logical indicator specifying if the `x` variable in the `formula` argument is already on the log scale. The default value is `TRUE`, thus, if `x` is given on a natural scale, `is_log` argument should be set to `FALSE`.

Arguments `lower_bound` and `upper_bound` are used for performing constrained optimization. They serve as the minimum and maximum values allowed for the model parameters. They are vectors of length equal to the number of parameters of the model specified by the `mean_function` argument. Values `-Inf` and `Inf` are allowed. The parameters for the 5-parameter generalized logistic function are listed in the following order: *α, β, η, ϕ, ν*. For the other models the order is preserved but some of the parameters are excluded. Obviously, values in `upper_bound` must be greater than or equal to the corresponding values in `lower_bound`.

`start` represents a vector of starting values for the parameters.

Finally, the `max_iter` argument sets the value for the maximum number of iterations in the optimization algorithm.

After the call to `drda()`, all the common functions expected for a model fit are available: `coef(), deviance(), logLik(), plot(), predict(), residuals(), sigma(), summary(), weights()`.

To evaluate the efficacy of the treatment it is also possible to compute the normalized area under or above the curve. The functions are respectively

~~~
nauc(drda_object, xlim = c(−10, 10), ylim = c(0, 1))
naac(drda_object, xlim = c(−10, 10), ylim = c(0, 1))
~~~

The two-element vector `xlim` defines the interval of integration, on the log-scale, with respect to `x`. The two-element vector `ylim` defines the theoretical minimum and maximum values for the response variable `y`. Therefore, `xlim` and `ylim` together define a rectangle that is partitioned into two regions by the dose-response curve. The normalized area under the curve (NAUC) is defined as the area of the “lower” rectangle region divided by the total area of the rectangle. The normalized area above the curve (NAAC) is simply its complement, i.e. 1 - NAUC.

When `xlim` and `ylim` are not explicitly chosen, the default values are set respectively to `c(−10, 10)` and `c(0, 1)`. The `xlim` default value was chosen on the basis of dose ranges that are commonly found in the literature, and made symmetric around zero so that NAUC and NAAC values are equal to 0.5 in the standard logistic model. In the majority of real applications the response variable `y` is usually a relative measure against a control treatment, therefore the default value for `ylim` is chosen to be `c(0, 1)`.

### 3.2. Usage examples

First of all, we load the package.

~~~
R> library(drda)
~~~

We then define an example dataset to demonstrate how to use **drda**.

~~~
R> dose <- rep(c(0.0001, 0.001, 0.01, 0.1, 1, 10, 100), each = 3)
R> relative_viability <- c(
+ 0.877362, 0.812841, 0.883113, 0.873494, 0.845769, 0.999422, 0.888961,
+ 0.735539, 0.842040, 0.518041, 0.519261, 0.501252, 0.253209, 0.083937,
+ 0.000719, 0.049249, 0.070804, 0.091425, 0.041096, 0.000012, 0.092564
+)
~~~

This example imitates an experiment where seven drug doses have been tested three times each. Relative viability measures have been obtained for each dose-replicate pair and, in this case, comprise 21 values in the (0, 1) interval. Note that any finite real number is accepted as a possible valid outcome.

#### Default fitting

The `drda()` function can be applied directly to the two variables via setting `is_log` to `FALSE`.

~~~
R> fit <- drda(relative_viability ~ dose, is_log = FALSE)
~~~

We can obtain exactly the same fitting using the log-doses and ignoring the `is_log` argument and storing the variables into a data frame.

~~~
R> log_dose <- log(dose)
R> test_data <- data.frame(d = dose, x = log_dose, y = relative_viability)
R> # the following calls are equivalent
R> fit <- drda(relative_viability ~ log_dose)
R> fit <- drda(y ~ d, data = test_data, is_log = FALSE)
R> fit <- drda(y ~ x, data = test_data)
~~~

To obtain summaries the user can apply the `summary()` function to the `fit` object.

|  | Min | 1Q | Median | 3Q | Max |
| --- | --- | --- | --- | --- | --- |
|  | -1.81632 | -0.45751 | -0.02341 | 0.20617 | 2.04357 |

Parameters:
|  | Estimate | Lower .95 | Upper .95 |
| --- | --- | --- | --- |
| Minimum | 0.05207 | -0.001967 | 0.106 |
| Maximum | 0.87914 | 0.828166 | 0.930 |
| Growth rate | -1.14335 | -1.683129 | -0.604 |
| Midpoint at | -2.11770 | -2.476042 | -1.759 |
| Residual std err. | 0.06541 | 0.042006 | 0.089 |

Residual standard error on 17 degrees of freedom
Log-likelihood: 29.688
AIC: -51.377
BIC: -47.199
Optimization algorithm converged in 20 iterations

The `summary()` function provides information about the Pearson residuals, parameters’ and residual standard error estimates, and their 95% confidence intervals. Together with the actual point estimate, the widths of confidence intervals are a good starting point for assessing the reliability of the model fit. The values of the log-likelihood function, AIC, and BIC are also provided. Finally, the `summary()` function warns the user if the algorithm converges and if so, in how many iterations.

Parameter estimates can be accessed using the `coef()` and `sigma()` functions, or by accessing them directly.

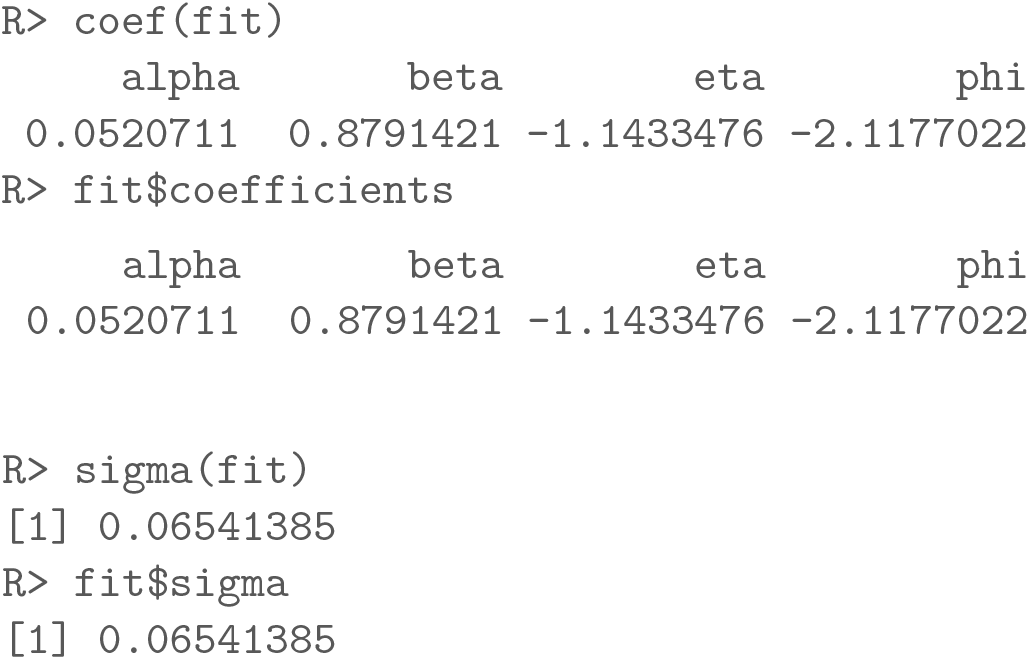

Since the model being fitted is (1), it is important to note that the the `coef()` function always returns the parameter *ϕ*, in this case the log-EC50, regardless of the scale in which `x` was passed to the function. The `summary()` function, however, will always print the estimate on the same scale as the original `x` variable.

Our `fit` object can be further explored with all the familiar functions expected for a model fit:

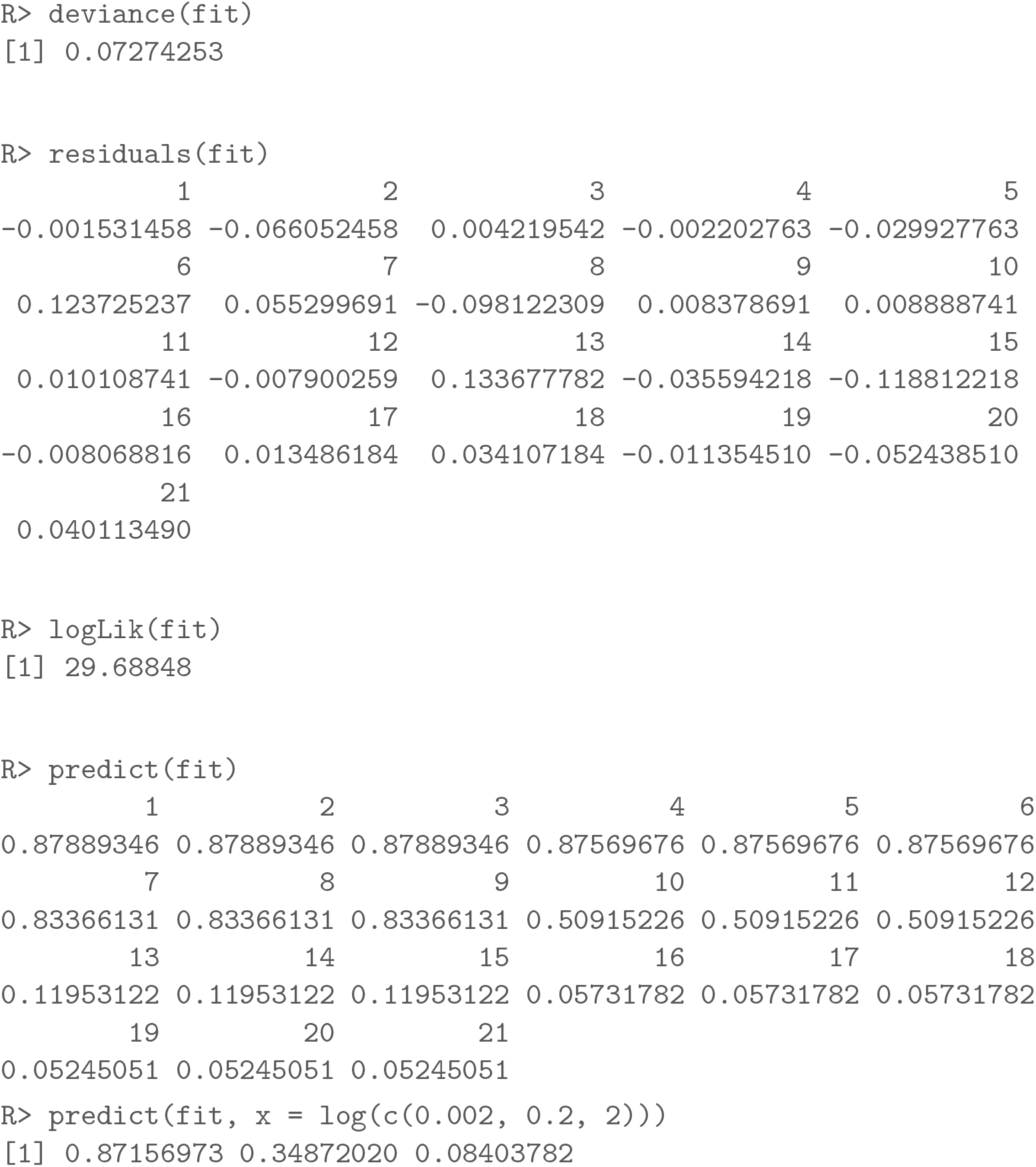

#### Model comparison and selection

The `anova()` function can be used to compare competing models within the same logistic family of models. The constant model, i.e. a flat horizontal line, is always included by default in the comparisons. When the model being fitted is not the 5-parameter logistic function, the latter is always included as the general reference model in the likelihood-ratio test.

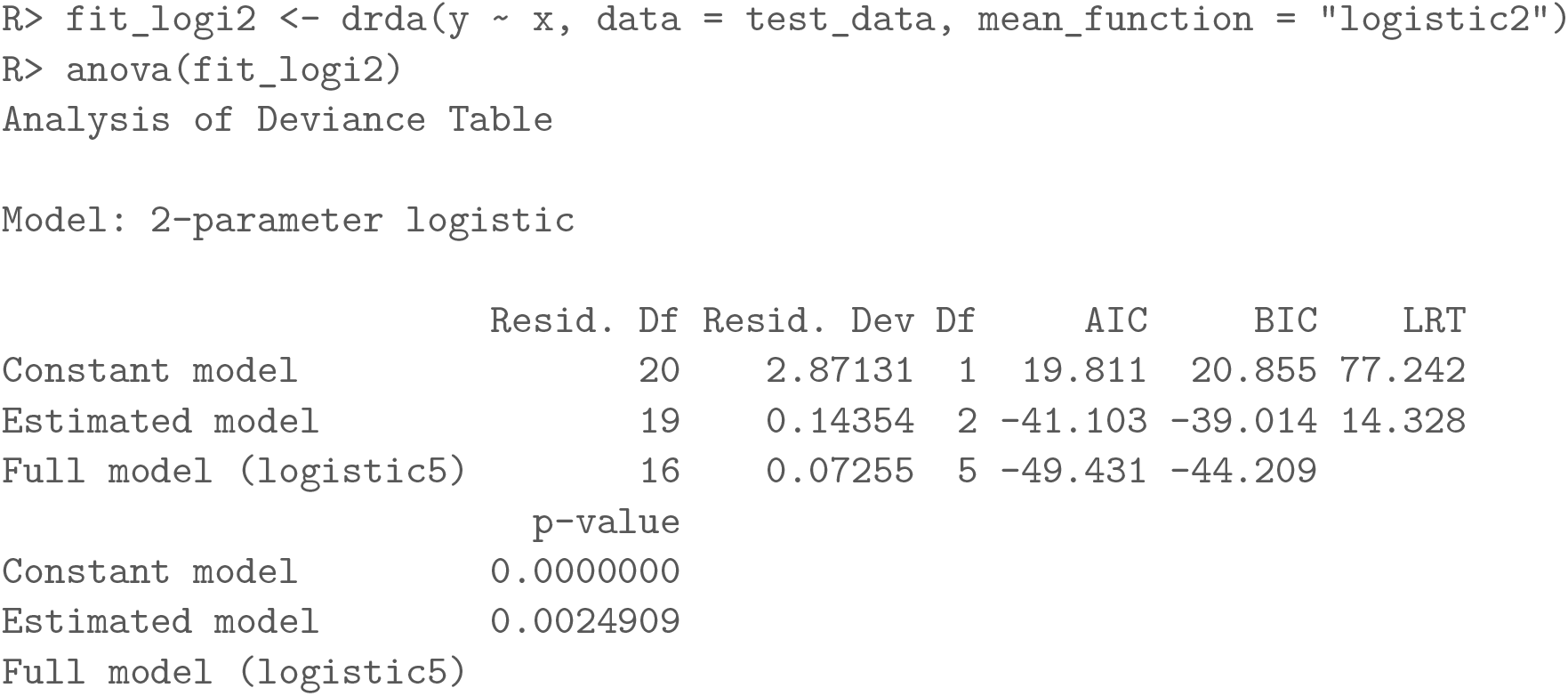

|  | Resid. Df | Resid. Dev | Df | AIC | BIC | LRT |
| --- | --- | --- | --- | --- | --- | --- |
| Constant model | 20 | 2.87131 | 1 | 19.811 | 20.855 | 77.242 |
| Estimated model | 19 | 0.14354 | 2 | -41.103 | -39.014 | 14.328 |
| Full model (logistic5) | 16 | 0.07255 | 5 | -49.431 | -44.209 |  |

p-value

|  |  |
| --- | --- |
| Constant model | 0.0000000 |
| Estimated model | 0.0024909 |
| Full model (logistic5) |  |

Note that the p-value refers here to the likelihood-ratio test with a $\chi^2$ -distribution asymptotic approximation. In this particular case we are testing the null hypothesis that our 2-parameter logistic function is equivalent, likelihood-wise, to the complete 5-parameter logistic function. The significant result indicates that the 2-parameter logistic function provides a worse fit for the observed data compared to a 5-parameter logistic function.

Note that the p-value refers here to the likelihood-ratio test with a *χ*^2^-distribution asymptotic approximation. In this particular case we are testing the null hypothesis that our 2-parameter logistic function is equivalent, likelihood-wise, to the complete 5-parameter logistic function. The significant result indicates that the 2-parameter logistic function provides a worse fit for the observed data compared to a 5-parameter logistic function.

|  | Resid. Df | Resid. Dev | Df | AIC | BIC | LRT | p-value |
| --- | --- | --- | --- | --- | --- | --- | --- |
| Model 1 | 20 | 2.87131 | 1 | 19.811 | 20.855 | 77.242 | 0.00000 |
| Model 2 | 19 | 0.14354 | 2 | -41.103 | -39.014 | 14.328 | 0.00077 |
| Model 3 | 17 | 0.07595 | 4 | -50.472 | -46.294 | 0.959 | 0.00000 |
| Model 4 | 17 | 0.07274 | 4 | -51.377 | -47.199 | 0.054 | 0.81569 |
| Model 5 | 16 | 0.07255 | 5 | -49.431 | -44.209 |  |  |

These results indicate the 4-parameter logistic function as the best fit for the data. Not only the model has the lowest AIC value, but the LRT is also not significant. Indeed, the data was generated from a 4-parameter logistic function with ***ψ*** = (0.02, 0.86, −1, −2) and *σ* = 0.05.

#### Weighted fitting

In case when not all of the observations should be utilized equally in the model, the weights argument can be provided to the `drda()` function. All the generic functions described above are also applicable to a weighted fit object.

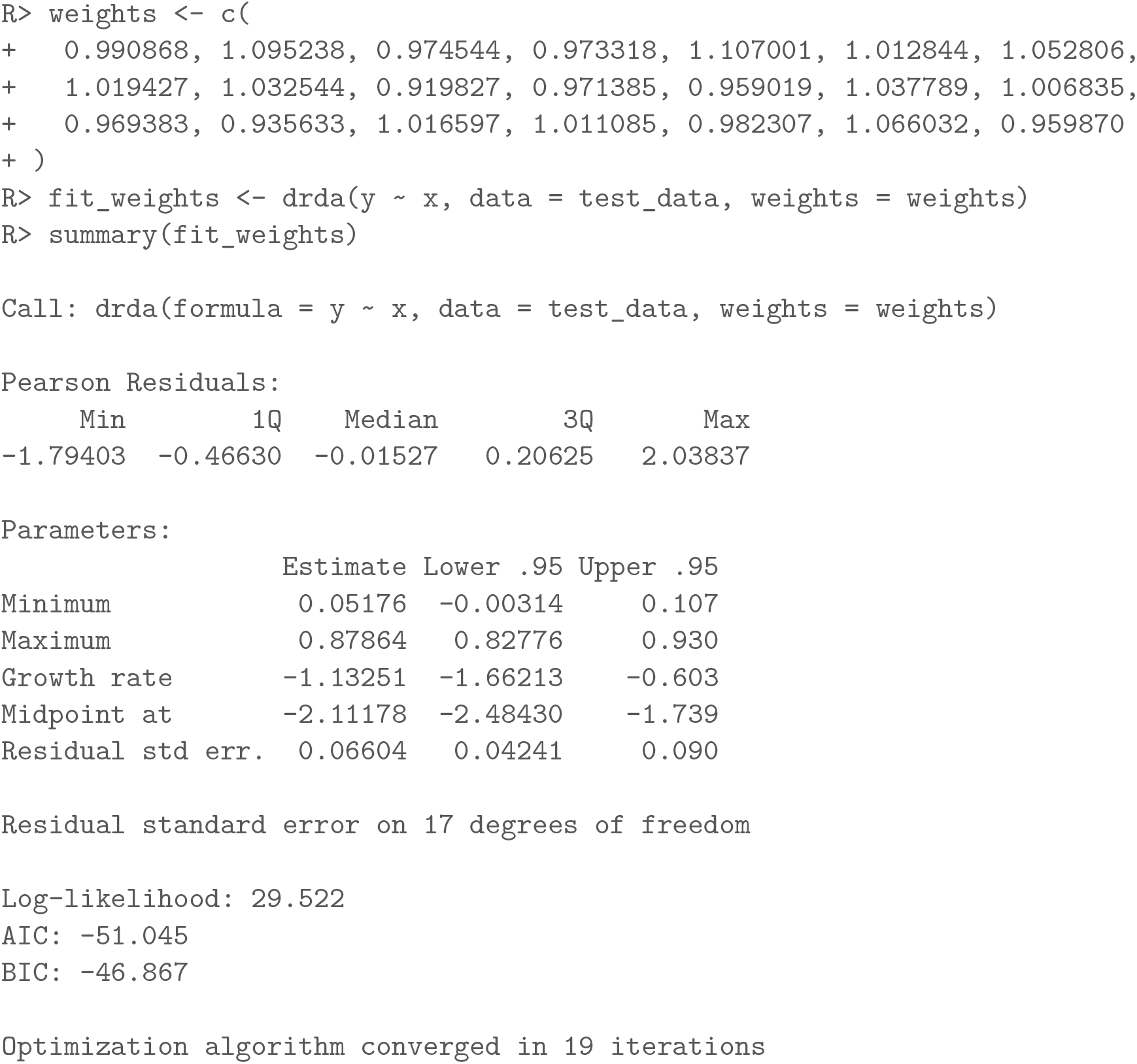
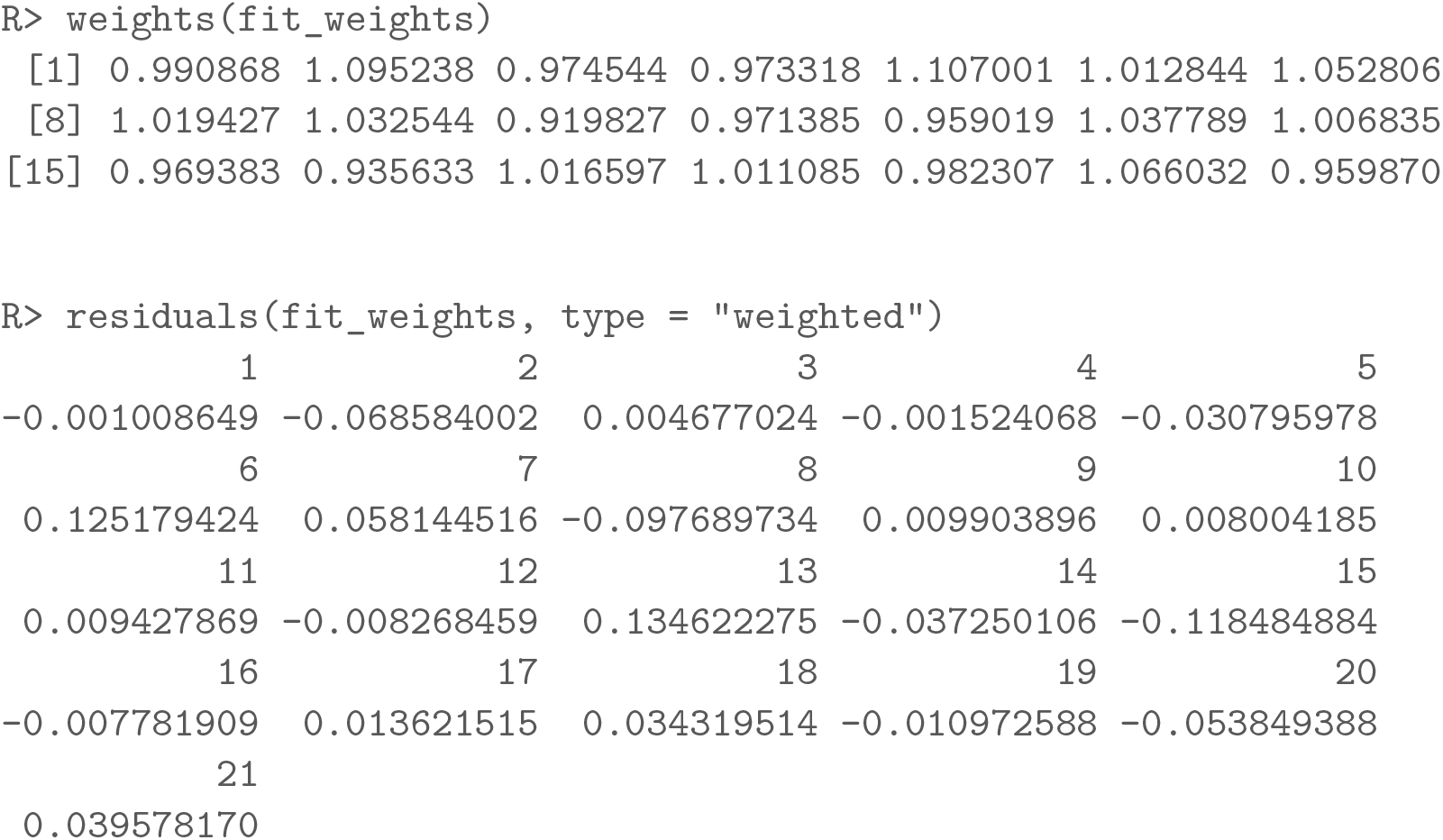

#### Constrained optimization

The `drda()` function allows the choice of admissible values for the parameters by setting the `lower_bound` and `upper_bound` arguments appropriately. Unconstrained parameters are set to `−Inf` and `Inf` respectively. While setting the constraints manually, one should be careful in choosing the values as the optimization problem might become very difficult to solve within a reasonable number of iterations.

In the next example the lower bound and upper bound parameters are fixed to 0 and 1 respectively, the growth rate is allowed to vary in [−5, 5], while the midpoint parameter is left unconstrained.

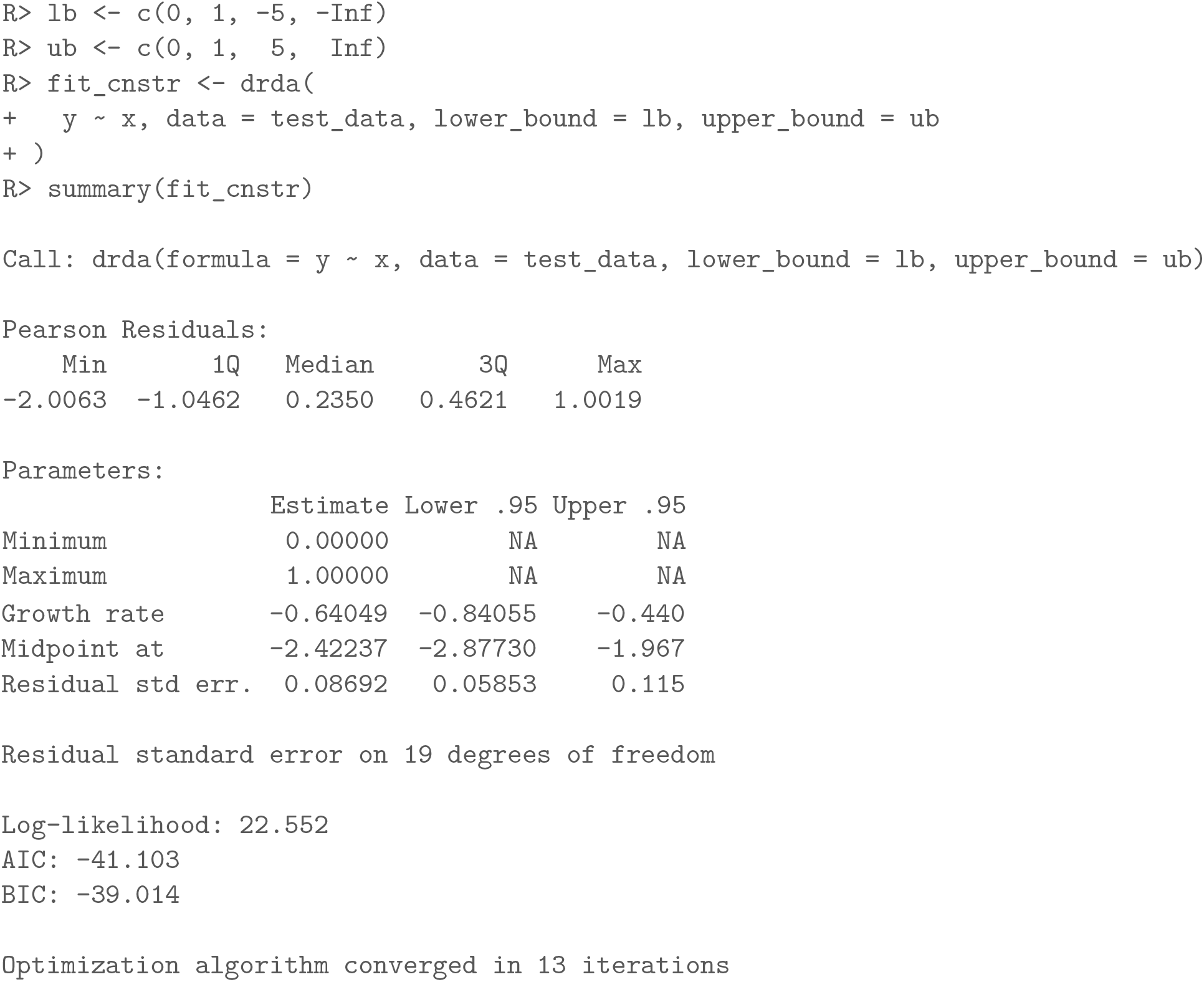

| Min | 1Q | Median | 3Q | Max |
| --- | --- | --- | --- | --- |
| -2.0063 | -1.0462 | 0.2350 | 0.4621 | 1.0019 |

Parameters:
|  | Estimate | Lower .95 | Upper .95 |
| --- | --- | --- | --- |
| Minimum | 0.00000 | NA | NA |
| Maximum | 1.00000 | NA | NA |
| Growth rate | -0.64049 | -0.84055 | -0.440 |
| Midpoint at | -2.42237 | -2.87730 | -1.967 |
| Residual std err. | 0.08692 | 0.05853 | 0.115 |

Finally, it is possible to provide an explicit starting point using the `start` argument or change the maximum number of iterations with the `max_iter` argument.

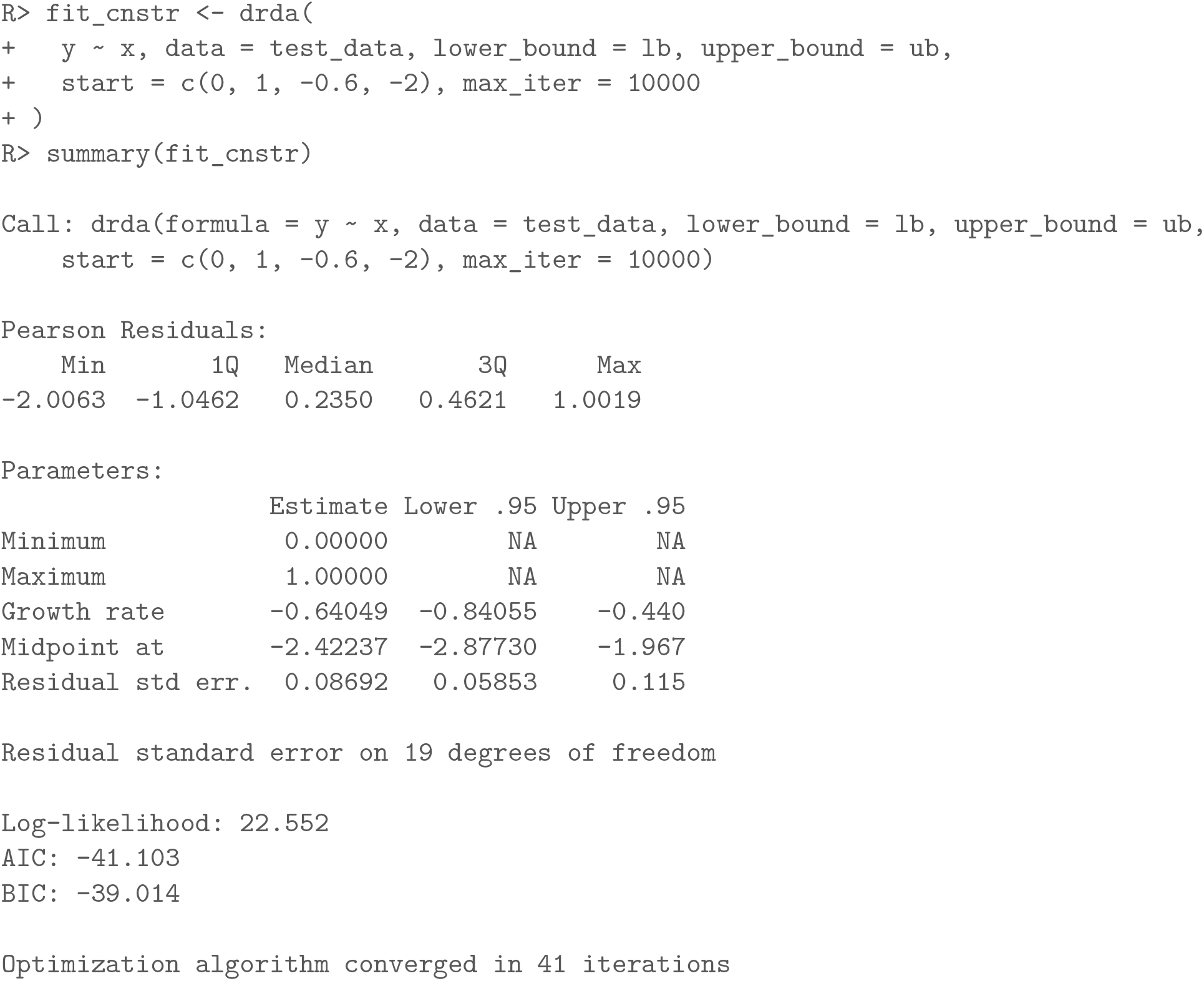

#### Basic plot functionality

As basic plot functionality, **drda** allows to plot the data used for fitting, the maximum likelihood curve and the approximate confidence intervals for the curve.

~~~
R> fit_logi5 <- drda(y ~ x, data = test_data, mean_function = “logistic5”)
R> plot(fit_logi5)
~~~

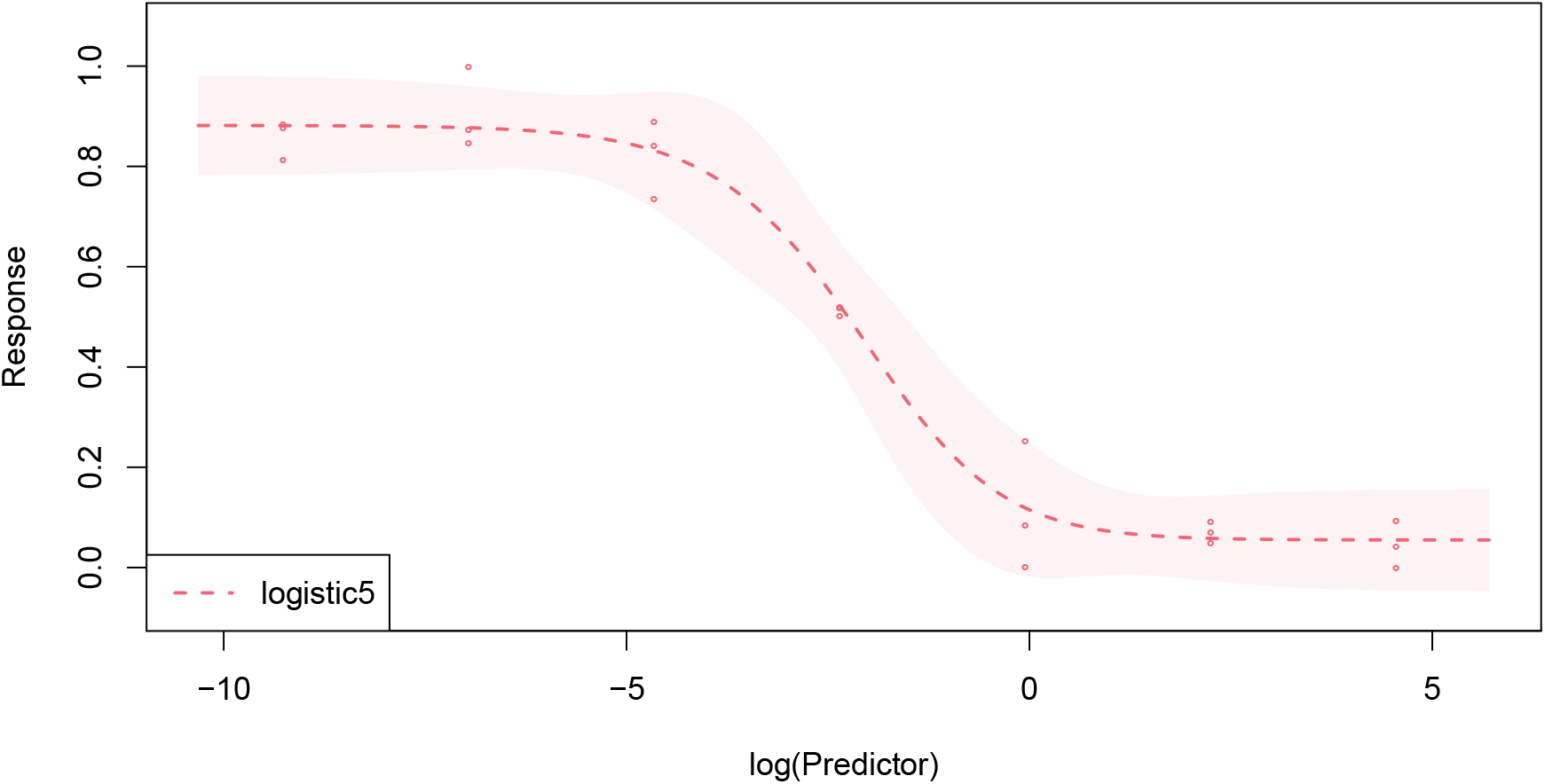

Alongside the common `plot()` arguments, it is possible to customize the plot by changing the scale of the x-axis with the argument `base` or the level of the confidence intervals with the `level` argument (default to 0.95). The available options for `base` are `‘e’, ‘2’`, and `‘10’`, with the default setting depending on the scale used for the `x` variable in the model `formula`. When the 2- or 4-parameter logistic functions are plotted, the *ϕ* parameter is also shown in the plot. It is also possible to plot any number of models within the same figure.

~~~
R> plot(
+ fit_logi2, fit_logi4, fit_gompe,
+ base = “10”, level = 0.9,
+ xlim = c(−10, 5), ylim = c(−0.1, 1.1),
+ xlab = “Dose”, ylab = “Relative viability”,
+ legend = c(“2-param logistic”, “4-param logistic”, “Gompertz”)
+)
~~~

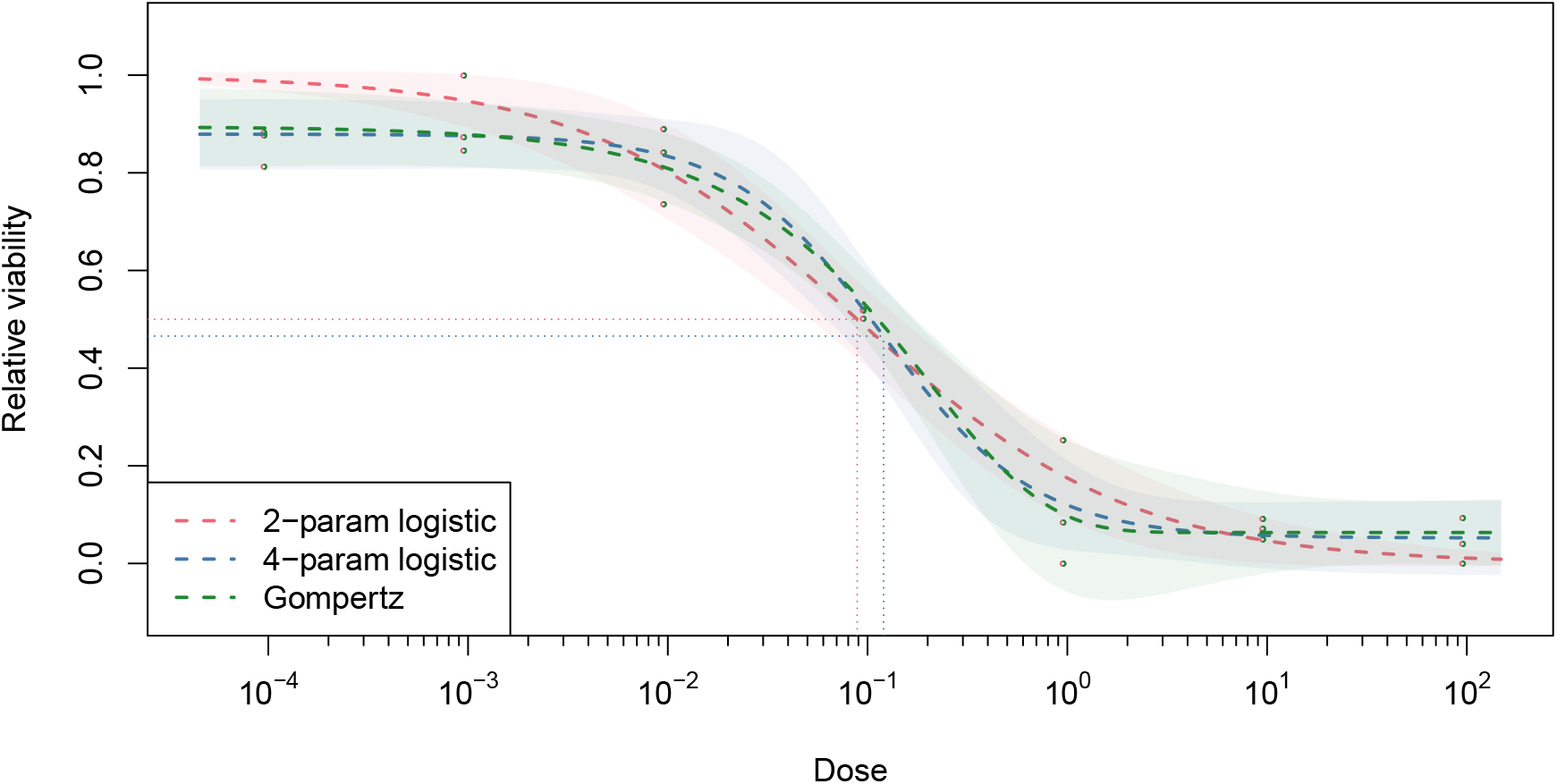

#### Area-based metrics

To obtain a measure of treatment efficacy, functions `nauc()` and `naac()` compute respectively the normalized area under the curve and above the curve. Since our example data refers to viability data, we use here the NAAC measure: the closer the value to 1 the better the treatment effect.

~~~
R> naac(fit_logi4)
[1] 0.6219635
~~~

To allow the values to be comparable between different compounds and/or studies, the function sets a hard constraint on both the `x` and `y` variables (see Section 3.1). However, the intervals can be easily changed if needed.

~~~
R> naac(fit_logi4, xlim = c(−2, 2), ylim = c(0.1, 0.9))
[1] 0.9062705
~~~

## 4. Benchmarking

We will now assess the performance and estimation accuracy of **drda** using a real large-scale drug sensitivity dataset downloaded from the Cancer Therapeutics Response Portal (CTRP) (Rees *et al*. 2016; Seashore-Ludlow *et al*. 2015; Basu *et al*. 2013). The data contains cell viability measures for 387130 cell line/drug pairs (887 unique cell lines, 545 unique drugs). The majority of experiments (79.3%) were performed for sixteen drug doses and no replicates, which is only one observation per dose. The relative viability measures span the (0.0019, 2.881) interval.

To choose reference values to compare our package to, we fitted the same model with the three packages - **DoseFinding**, **drc**, and **nplr**. As a control variable for the comparison, we chose the 4-parameter logistic model in all packages, and the arguments of each package core function were set to produce results that are as similar as possible. For `drm()` from package **drc**, we selected the 4-parameter logistic model with `fct = L.4()` and fixed the maximum number of iterations to 10000, similarly to `drda()`. For `nplr()` from package **nplr**, we changed useLog to `FALSE` and set `LPweight` to 0 in order to perform the ordinary least squares method. We fixed `npars` to four for the 4-parameter logistic model. For `fitMod()` from package **DoseFinding** we chose the 4-parameter logistic model by setting model = `“sigEmax”` (see Section 2.1). Since the `fitMod()` function requires the user to set constraints on the nonlinear parameters, we used the default value `bnds = defBnds(max(dose))$sigEmax`.

For each cell line-drug-package triple we fitted the 4-parameter logistic function one hundred times with function `benchmark()` from `R` package **rbenchmark** (Kusnierczyk 2012) and recorded the parameter estimates, the residual standard error, the residual sum of squares (RSS), convergence status, and the elapsed time of the function.

Since all packages are solving the same optimization problem (2), i.e. minimization of the residual sum of squares, we considered for each cell line-drug pair the global optimum to be the fit with the lowest RSS value among the four packages. We define the absolute relative error of package *k* as

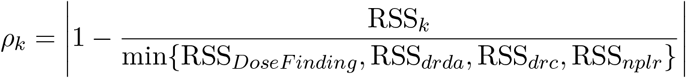

For real applications, small absolute relative errors (here we set the threshold to 0.01) can be considered equivalent to zero. Results are shown in Table 1.

**Table 1:** Summary statistics of benchmarking results for package **drda**.

| Statistic | Relative error |  |  |  |
| --- | --- | --- | --- | --- |
|  | **drda** | **DoseFinding** | **drc** | **nplr** |
| Minimum | 0.0000 | 0.000 | 0.00000 | 0.000 |
| First quartile | 0.0000 | 0.000 | 0.00003 | 0.000 |
| Median | 0.0000 | 0.000 | 0.00352 | 0.109 |
| Mean | $1.39 \cdot 10^{-5}$ | 0.778 | 0.06400 | 4.556 |
| Third quartile | 0.0000 | 0.020 | 0.04290 | 2.329 |
| Maximum | 0.2493 | 12157.546 | 218.32983 | 8338.883 |

Overall, **drda** is flagged as the absolute best fit in 90.81% of cases. When we only consider the cases for which |*ρ_k_*| ≤ 0.01, the percentage raises to 99.96% (70.21% for **DoseFinding**, 59.98% for **drc**, and 43.65% for **nplr**). When compared directly against the other packages, **drda** outperforms **DoseFinding** in 29.78% of the cases (worse for 0.033%), **drc** in 39.99% of cases (worse for 0.004%), and **nplr** in 56.34% of the cases (worse for 0.016%).

The results show that **drda** provides more accurate, and thus more reliable, estimates of the dose-response relationship. The higher accuracy comes obviously at a computational cost, asmore steps are usually needed for exploring the parameter space. Our data analysis reveals that `fitMOD()` and `nplr()` are the fastest functions to complete the fit. It took them less than a second to converge 95% of the times (mean of 0.62s and median of 0.61s for `fitMOD()`; mean of 0.91s and median of 0.95s for `nplr()`). On average **drda** found the global optimum (or a very close solution) in 14.45 seconds (median of 9.6s). For completeness, `drm()` had an average of 9.87 seconds and a median of 3.27 seconds.

## 5. Summary and discussion

In this paper, we have introduced the **drda** package, aimed at evaluating dose-response relationship to advance our understanding of biological processes or pharmacological safety. These types of experiments are of high importance in drug discovery, as they establish an essential step for subsequent therapeutic advances. An appropriate interpretation of the experimental data is grounded on a reliable estimation of the dose-response relationship. Therefore, it is imperative to provide advanced optimization methods that allow more accurate estimation of dose-response parameters, and the assessment of their statistical significance.

One of the main limitations of most optimization procedures is their convergence to local solutions. The basic quasi-Newton methods applied to logistic curve fitting are sensitive to the selection of a starting point and to cases when data is non-informative. Our package effectively overcomes the convergence problem as we implement a Newton method with a trust region to achieve global convergence and improve it further with a double-step starting point initialization. The **drda** optimization routine also relies on analytical gradient and Hessian to avoid numerical approximations. The package allows a user to further evaluate the model fitness further via the assessment of confidence intervals of the estimates, model comparisons, and advanced plot options.

We have compared our package with the three state-of-the-art packages - **DoseFinding**, **drc**, and **nplr**. Using a large-scale drug screening dataset, we have shown that **drda** has clearly outperformed the other three packages in terms of accuracy. Despite the fact that our package is on average slower than the other three packages, its gain in accuracy is a favorable compromise. For most, if not all, experimental applications, accuracy has a higher priority. The package is currently completely implemented in base `R`, therefore there are still many opportunities for improving its performance, by, for example, refactoring core critical functions in `C` or improving further the algorithm initialization. If a researcher is looking for a package providing improved accuracy at a relatively low speed-cost, **drda** might provide a viable option. The package can be downloaded from https://github.com/albertopessia/drda.

## Acknowledgments

We thank CSC, the Finnish IT center for science, for the computational resources used to perform the simulations.

Research is supported by the European Research Council (ERC) starting grant, No 716063 (DrugComb: Informatics approaches for the rational selection of personalized cancer drug combinations).

